# Genetic diversity of clinical *Bordetella pertussis* ST2 strains in comparison with vaccine reference strains of India

**DOI:** 10.1101/573543

**Authors:** Naresh Chand Sharma, Shalini Anandan, Naveen Kumar Devanga Ragupathi, Dhiviya Prabaa Muthuirulandi Sethuvel, Karthick Vasudevan, Dhirendra Kumar, Sushil Kumar Gupta, Lucky Sangal, Vijayalakshmi Murugesan, Balaji Veeraraghavan

**Affiliations:** Maharishi Valmiki Infectious Diseases Hospital, New Delhi – 110009, India; Department of Clinical Microbiology, Christian Medical College, Vellore – 632 004, India; World Health Organization, Country Office, New Delhi – 110029, India

**Keywords:** *Bordetella*, *ptx*, IS*481*, genome reduction, ST2

## Abstract

Pertussis is a highly contagious disease of the respiratory tract caused by *Bordetella pertussis*, a bacteria that lives in the mouth, nose, and throat. Current study reports the highly accurate complete genomes of two clinical *B. pertussis* strains from India for the first time. The analysis revealed insertional elements flanked by IS*481*, which has been previously regarded as the important component for bacterial evolution. The two *B. pertussis* clinical strains exhibited diversity through genome degradation when compared to whole-cell pertussis vaccine reference strains of India. These isolates harboured multiple genetic virulence traits and toxin subunits, which belonged to sequence type ST2. The genome information of Indian clinical *B. pertussis* strains will serve as a baseline data to decipher more information on the genome evolution, virulence factors and their role in pathogenesis for effective vaccine strategies.

## Introduction

Pertussis, caused by *Bordetella pertussis,* is a highly contagious respiratory infection, characterized by severe episodes of coughing and a prolonged convalescent period when the patient can transmit the disease (1). Although large scale vaccination reduced the incidence of the disease, recent trends suggest a re-emergence of this disease particularly among the adolescent and young adult population in the developed countries. Genome data of *B. pertussis* is rare due to the complexity of culturing *B. pertussis* from clinical samples. Understanding the molecular composition of these isolates is essential to obtain the molecular epidemiological information that will be helpful in public heath surveillance. Two *B. pertussis* isolates (BPD1, BPD2) identified from nasopharyngeal swabs of paediatric patients belonging to Uttar Pradesh, India were confirmed by culture and real-time PCR (2).

## Materials and Methods

### Strain Isolation and Characterization

Bacterial strains BPD1 and BPD2 were isolated from nasopharyngeal swabs of paediatric patients. Swabs were plated in charcoal blood agar and incubated at 37 °C with CO_2_ for 48 hours. Isolates were confirmed by standard biochemical tests and real-time PCR for the targets IS*481* and *ptxS1* genes.

### Genome sequencing

#### Short read sequencing and assembly

Genomic DNA of the *B. pertussis* isolates were extracted using QIAamp DNA Mini Kit (QIAGEN, Hilden, Germany). Whole genome sequencing (WGS) was performed in IonTorrent™ Personal Genome Machine™ (PGM) (Life Technologies, Carlsbad, CA) with 400-bp read chemistry as per manufacturer’s instructions. Raw reads were assembled *de-novo* using Assembler SPAdes v.5.0.0.0 embedded in Torrent Suite Server v.5.0.3.

#### Long read sequencing and assembly

Library preparation and sequencing of the *B. pertussis* isolates was done using SQK-LSK108 Kit R9 version (Oxford Nanopore Technologies, Oxford, UK) using 1D sequencing method according to manufacturer’s protocol. Sequencing of the isolates was performed using FLO-MIN106 R9 flow cell in MinION Mk 1B sequencer. To perform sequencing, MinKNOW software ver. 1.15.1 (Oxford Nanopore Technologies, Oxford, UK) was used in a Windows platform and raw data (fast5 files) were obtained. The Fast5 files were basecalled with Albacore 2.0.1 (https://nanoporetech.com/about-us/news/new-basecaller-now-performs-raw-basecalling-improved-sequencing-accuracy). Error correction and genome assembly was performed using Canu 1.7 (Koren et al. 2017). The obtained contigs were polished with Nanopolish 0.10.1 (https://github.com/jts/nanopolish) after *de novo* assembly.

#### Hybrid assembly using IonTorrent and MinION reads

Hybrid assembly using both IonTorrent and MinION reads were performed to increase the accuracy and completeness of genome. Unicycler (v0.4.6) was used for generating hybrid assemblies (Wick et al. 2017). Further, the reads were polished with multiple rounds of Pilon (Walker et al. 2014) (4) to reduce the base level errors. The assembly statistics and average nucleotide identity of different assemblies were evaluated using Quast (Gurevich et al. 2013).

### Genome annotation and MLST analysis

Annotation of the sequences were done using PATRIC, the bacterial bioinformatics database and analysis resource (http://www.patricbrc.org) (5), and NCBI Prokaryotic Genomes Automatic Annotation Pipeline (PGAAP, http://www.ncbi.nlm.nih.gov/genomes/static/Pipeline.html). MLST 1.8 (MultiLocus Sequence Typing) tool was employed for sequence type analysis (https://cge.cbs.dtu.dk//services/MLST/) (Larsen et al. 2012).

## Results and Discussion

### Genome length, CDS and ST types

Hybrid assemblies returned with 203X and 195X coverage for BPD1 and BPD2 isolates respectively for the complete genomes in MinION platform. The completed BPD1 genome had a sequence length of 4,126,211 bp with 3941 CDS, 3 rRNA and 50 tRNA, and isolate BPD2 had 4,104,911 bp with 3921 CDS, 3 rRNA and 51 tRNA (https://www.patricbrc.org). The whole genome MLST using MLST 1.8 tool (https://cge.cbs.dtu.dk//services/MLST/) (6) revealed the sequence type (ST) of both isolates to be ST2 belonging to CC2, previously reported to be unique to Africa (7).

### Insertional elements observed in vaccine reference strains

BPD1 had included ∼128 Kb repeat insertion flanked by copies of IS*481* in single copy (Table 1), while, BPD2 had an insertion of ∼150 Kb length. However, these isolates lacked other repeat regions that were observed in the vaccine reference strains 6229 (CP017404) and 25525 (CP017405) used for production of whole-cell pertussis vaccine in India belonging to ST2 (8). Comparison of the repeat region observed in BPD1 with 25525 reference strain using Easyfig v2.2.3 showed the similarity between the two regions with internally inverted repeat regions (Figure 1). Whereas, the comparison of the BDP2 with 25525 genome exhibits the presence of a repeat region different than in the vaccine reference strains (Figure 2). These repeat regions mainly carry flagellar genes involved in pathogenesis of *B. pertussis*. Such transposable DNA elements were regarded as the potent force in the evolution of bacteria (Stibitz, 1998).

**Table 1:**
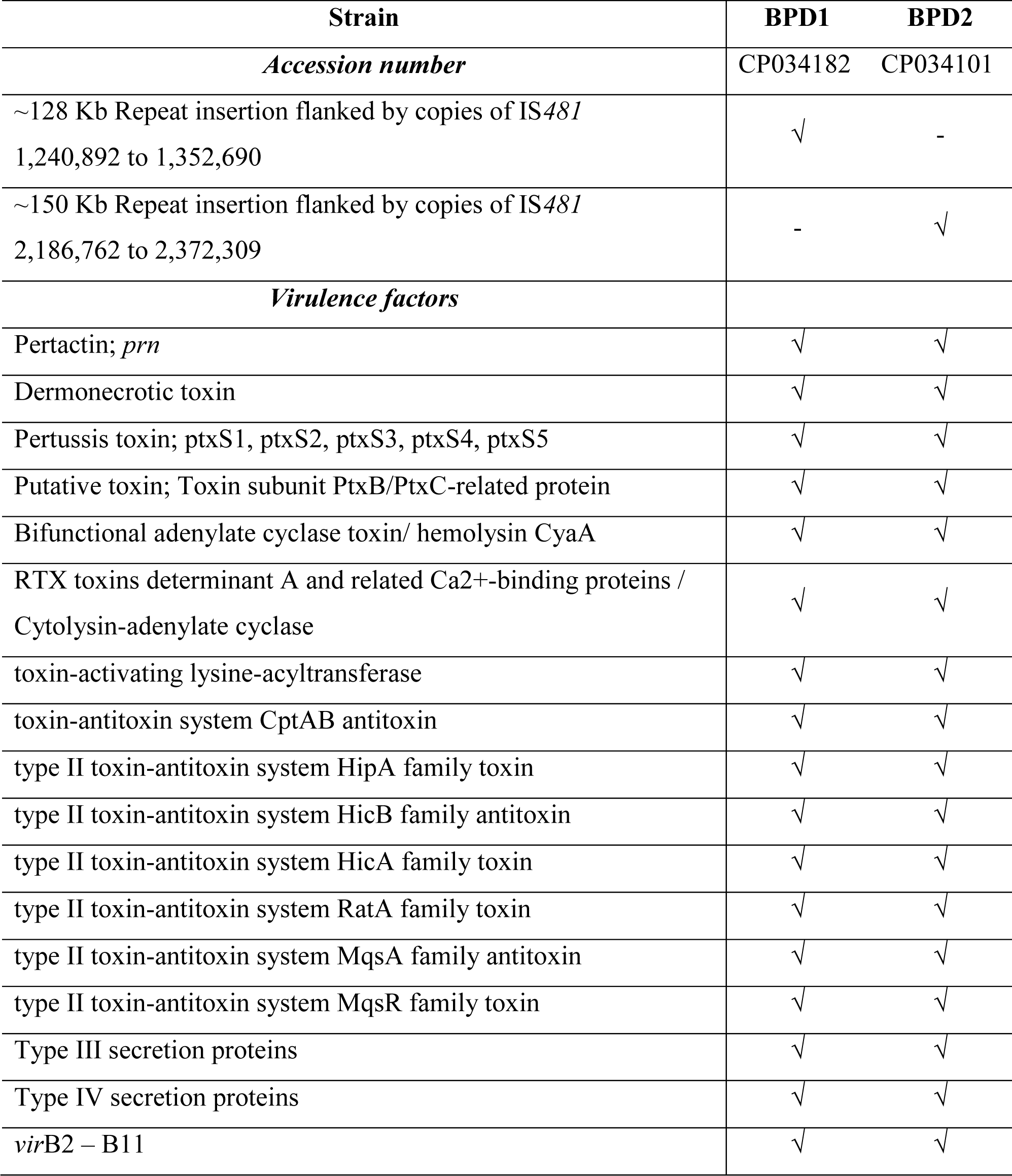
Virulence genome characteristics of clinical *B. pertussis* from India

| Strain | BPD1 | BPD2 |
| --- | --- | --- |
| *Accession number* | CP034182 | CP034101 |
| ~128 Kb Repeat insertion flanked by copies of IS481<br>1,240,892 to 1,352,690 | √ | - |
| ~150 Kb Repeat insertion flanked by copies of IS481<br>2,186,762 to 2,372,309 | - | √ |
| *Virulence factors* |  |  |
| Pertactin; *prn* | √ | √ |
| Dermonecrotic toxin | √ | √ |
| Pertussis toxin; ptxS1, ptxS2, ptxS3, ptxS4, ptxS5 | √ | √ |
| Putative toxin; Toxin subunit PtxB/PtxC-related protein | √ | √ |
| Bifunctional adenylate cyclase toxin/ hemolysin CyaA | √ | √ |
| RTX toxins determinant A and related Ca<sup>2+</sup>-binding proteins /<br>Cytolysin-adenylate cyclase | √ | √ |
| toxin-activating lysine-acyltransferase | √ | √ |
| toxin-antitoxin system CptAB antitoxin | √ | √ |
| type II toxin-antitoxin system HipA family toxin | √ | √ |
| type II toxin-antitoxin system HicB family antitoxin | √ | √ |
| type II toxin-antitoxin system HicA family toxin | √ | √ |
| type II toxin-antitoxin system RatA family toxin | √ | √ |
| type II toxin-antitoxin system MqsA family antitoxin | √ | √ |
| type II toxin-antitoxin system MqsR family toxin | √ | √ |
| Type III secretion proteins | √ | √ |
| Type IV secretion proteins | √ | √ |
| *virB2* – B11 | √ | √ |

**Figure 1:**
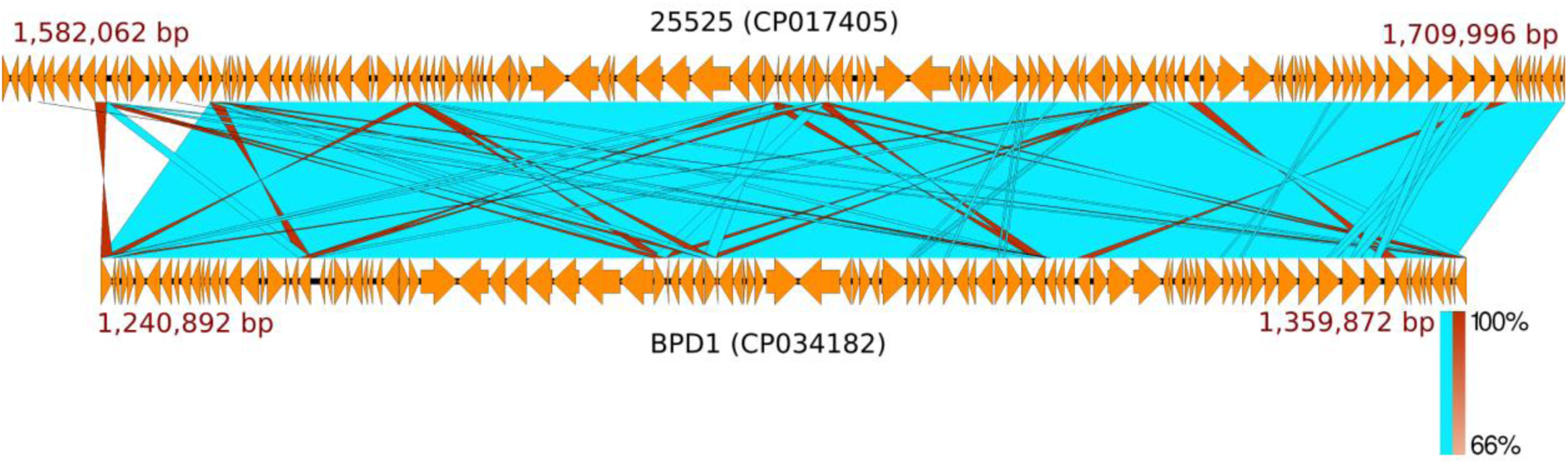
Representation of similarity and internal inverted repeats in comparison of repeat region flanked by IS*481* from BPD1 (CP034182) and the vaccine reference strain 25525 (CP017405)

**Figure 2:**
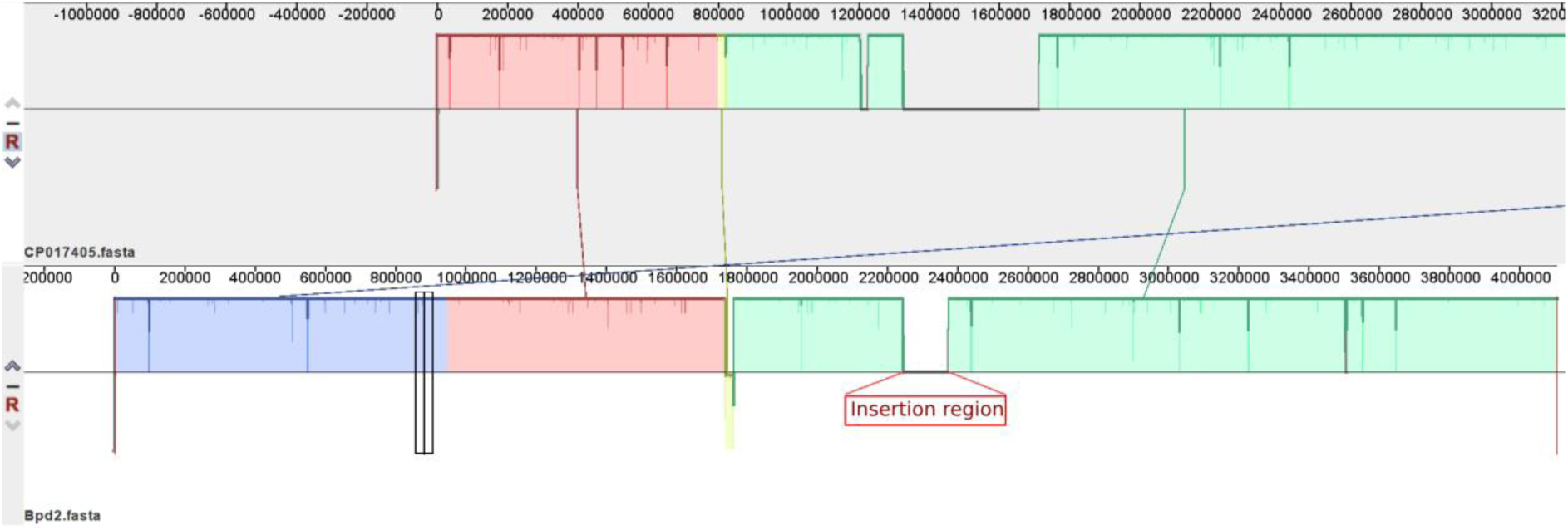
Comparison of BPD2 (CP034101) and 25525 (CP017405) revealing the presence of a repeat region in BPD2 different than in the vaccine strain.

Both, 6229 and 25525 were reported to be more closely related to the clinical strains than the other reference strains 134 (CP017402) and 509 (CP017403) (8). Homologous recombination between copies of IS*481* has been attributed to genome reduction in *B. pertussis* which also suggests possible genome expansion by the same mechanism. Similarly, BPD1 and BPD2 have undergone genome reduction due to IS*481* comparable to vaccine strains 6229 and 25525. The lesser number of structural genes adds up to the potential of *B. pertussis* to be more virulent as it reduces the number of targets that are available for recognition by the human immune system (9).

### Toxin and other virulence genes

Both BPD1 and BPD2 isolates had *ptx*S toxin genes with all five subunits. In addition, the isolates harboured pertactin adhesion (*prn*) gene, dermonecrotic, hemolysin and cytolysin toxins, and genes involved in type II toxin-antitoxin systems. Other genes identified include virulence genes such as *vir*B2, B3, B6, B7, B8, B9, B10 and B11 (Table 1).

### Pathogen identities

Moreover, the pathogenFinder 1.1 (https://cge.cbs.dtu.dk/services/PathogenFinder/) (10) for BPD1 proteome families matched with 10 non-pathogenic families and 3 pathogenic families. This showed sequence similarity of 88%, 85.9% and 87.18% to the pathogenic families of *Burkholderia cenocepacia, Burkholderia mallei* and *Neisseria gonorrhoeae* respectively. Similar results were observed for BPD2.

These genome information of *B. pertussis* isolates from India in comparison with vaccine reference strains will help to decipher more information on the genome evolution, virulence factors and their role in pathogenesis for effective vaccine strategies.

## Acknowledgements

The authors thank the funding agency, WHO Country Office for India for supporting this study. The research work has been approved by the Institutional Review Board of Christian Medical College, Vellore at the meeting conducted on 28/10/2016 (IRB Min No: 9706).

